# Expansion of DNA-Encoded Library Hits Using Generative Chemistry and Ultra-Large Compound Catalogs

**DOI:** 10.1101/2025.09.30.679600

**Authors:** Brandon Novy, Shu-Hang Lin, Devan J. Shell, Travis Maxfield, Eric M. Merten, Ivanna Zhilinskaya, James Wellnitz, Shiva K. R. Guduru, P. Brian Hardy, Kenneth H. Pearce, Konstantin I. Popov

**Affiliations:** UNC Eshelman School of Pharmacy, Center for Integrative Chemical Biology and Drug Discovery, Division of Chemical Biology and Medicinal Chemistry, University of North Carolina at Chapel Hill, Chapel Hill, NC 27599, USA; Lineberger Comprehensive Cancer Center, University of North Carolina at Chapel Hill, Chapel Hill, NC 27599, USA

## Abstract

DNA-encoded libraries (DELs) are powerful tools for initial hit identification, yet the combinatorial chemistries and building block choices used in their construction can restrict chemical space coverage and hit drug-likeness, limiting efficient hit expansion. Generative artificial intelligence (AI), by contrast, can in principle explore drug-like chemical space around any given compound, but it often struggles with the synthesizability of generated molecules and requires a set of validated hits to initiate exploration. Here, we present a synergistic methodology that overcomes these mutual limitations by leveraging experimentally validated DEL data to initialize and bias an AI-powered virtual screening pipeline, expanding initial DEL hits with both *de novo* and purchasable compounds from ultra-large chemical libraries. Using this approach, we identified novel and commercially available hits from the Enamine REAL Space for the chromatin reader protein 53BP1 and validated them in a time-resolved fluorescence resonance energy transfer (TR-FRET) displacement assay. Three compounds demonstrated TR-FRET IC50 values ≤50 µM, while 11 exhibited IC50 values ≤100 µM. Critically, the AI-nominated hits exhibited greater chemical diversity, improved drug-likeness, and were readily purchasable off-the-shelf compared to compounds from the initial DEL selection. This work demonstrates a streamlined platform in which empirical DEL data and generative chemistry models are combined to enable rapid hit expansion from initially screened libraries into diverse, commercially available chemical matter.

## INTRODUCTION

High-throughput screening (HTS) has long been a cornerstone of drug discovery, enabling the exploration of diverse or targeted chemical libraries ranging from thousands to millions of compounds.^1,2^ However, it often demands substantial resources and can be quite costly for academic labs and smaller drug discovery companies.^3^ To overcome the limitations of traditional HTS, combinatorial chemistry introduced methods for generating large libraries of small molecules, and with the advent of next-generation sequencing (NGS), DNA-encoded libraries (DELs) emerged as a powerful and resource-efficient alternative. By pooling diverse chemical building blocks tagged with unique DNA sequences, it is now routine to create and screen libraries with billions of compounds.^4^ NGS is used to read and decode DNA barcodes, track enrichment of chemical moieties in bound compounds, streamlining hit discovery. Numerous successes in DEL screening for hit discovery have been reported across various protein classes in both industry and academia.^5–10^ This technology has become integral to many pharmaceutical companies, demonstrating equal or greater success in efficient hit discovery compared to traditional HTS.^11,12^

Although the primary appeal of DELs is their ability to screen hundreds of millions to billions of molecules, smaller focused libraries targeting specific protein classes have also proven highly effective.^5,13^ This approach is supported by findings that library performance can decline with size; studies have reported decreasing signal-to-noise ratios and increasing false positive rates as library size increases.^14,15^ Our group, in a previous work, reported an in-house focused DEL containing around 58K members, UNCDEL003 (DEL3), that specifically designed to target methyl-lysine (Kme) reader domains.^16^ A key challenge with such focused libraries, given their smaller size compared to diversity-based DELs, is the need for methods that can explore chemical space beyond the confines of the original library.

To bridge this gap and bypass the resource-intensive bottleneck of off-DNA hit resynthesis, computational strategies to extrapolate DEL selection data into a larger chemical space have gained significant traction. Recently, McCloskey et al. demonstrated an effective approach using aggregated DEL screening data to train machine learning (ML) models for hit nomination from commercial libraries.^17^ They developed graph convolutional neural networks and a Random Forest model based on DEL selection data for three distinct targets, enabling virtual screening of commercial compounds. This approach identified structurally novel hits for all three proteins, highlighting the potential of DEL data to train ML models for discovering unique small molecule binders.^18^ These methods are well established for accelerating virtual screening across diverse chemical libraries, but their predictive capacity is constrained by the quality and scope of the training data. As a result, supervised learning models trained on DEL data may have difficulty generalizing to novel regions of drug-like chemical space.^19^

Recent advancements in computer-aided drug discovery (CADD) have enabled both traditional and AI-powered virtual screening (VS) of billion-size chemical libraries, making them comparable to DNA-encoded libraries (DELs) in the scale of chemical space explored for hit identification.^20,21^ Notably, successful structure-based VS provides a protein-ligand complex model, aiding downstream hit evaluation and optimization.^22^ We recently developed HIDDEN GEM, a structure-based virtual screening (VS) workflow that integrates molecular docking, biased generative AI, and efficient similarity searching to identify hits from ultra-large chemical libraries.^23^ Incorporating a generative step using SMILES-based language models pretrained on ChEMBL^24^ and biased by molecular docking proved crucial for navigating vast chemical space and identifying virtual hits. The workflow requires knowledge of the protein binding site and can be initialized with either a million-size diverse library or a smaller focused set if prior binders are known. HIDDEN GEM efficiently navigates ultra-large chemical libraries, identifying diverse high-scoring virtual hits with improved docking scores relative to the initialization set. The workflow can be iteratively repeated to further focus on a particular scaffold or increase hit diversity, with the output of each run used to initialize the subsequent one.

In this work, we test the hypothesis that initializing a structure-based generative AI workflow with validated DEL screening data can rapidly identify novel, potent, and more drug-like compounds from an ultra-large virtual library, outperforming traditional hit-picking strategies. To do so, we present a DEL-driven, integrated virtual screening workflow that combines the HIDDEN GEM platform with the multi-billion-compound Enamine REAL Space (**Figure 1**). Using data from our recently published DEL screens targeting the tandem Tudor domain (TTD) of 53BP1,^8,16,25^ we biased our generative structure-guided ML model to identify novel, commercially available compounds from the Enamine REAL Space. All purchased and off-DNA synthesized compounds were experimentally tested using a developed time-resolved fluorescence resonance energy transfer (TR-FRET) displacement assay. This workflow provides a rapid, cost-effective strategy to accelerate hit discovery using focused DELs and to expand beyond the screened chemical space, which is particularly important for smaller-sized DELs.

**Figure 1.**
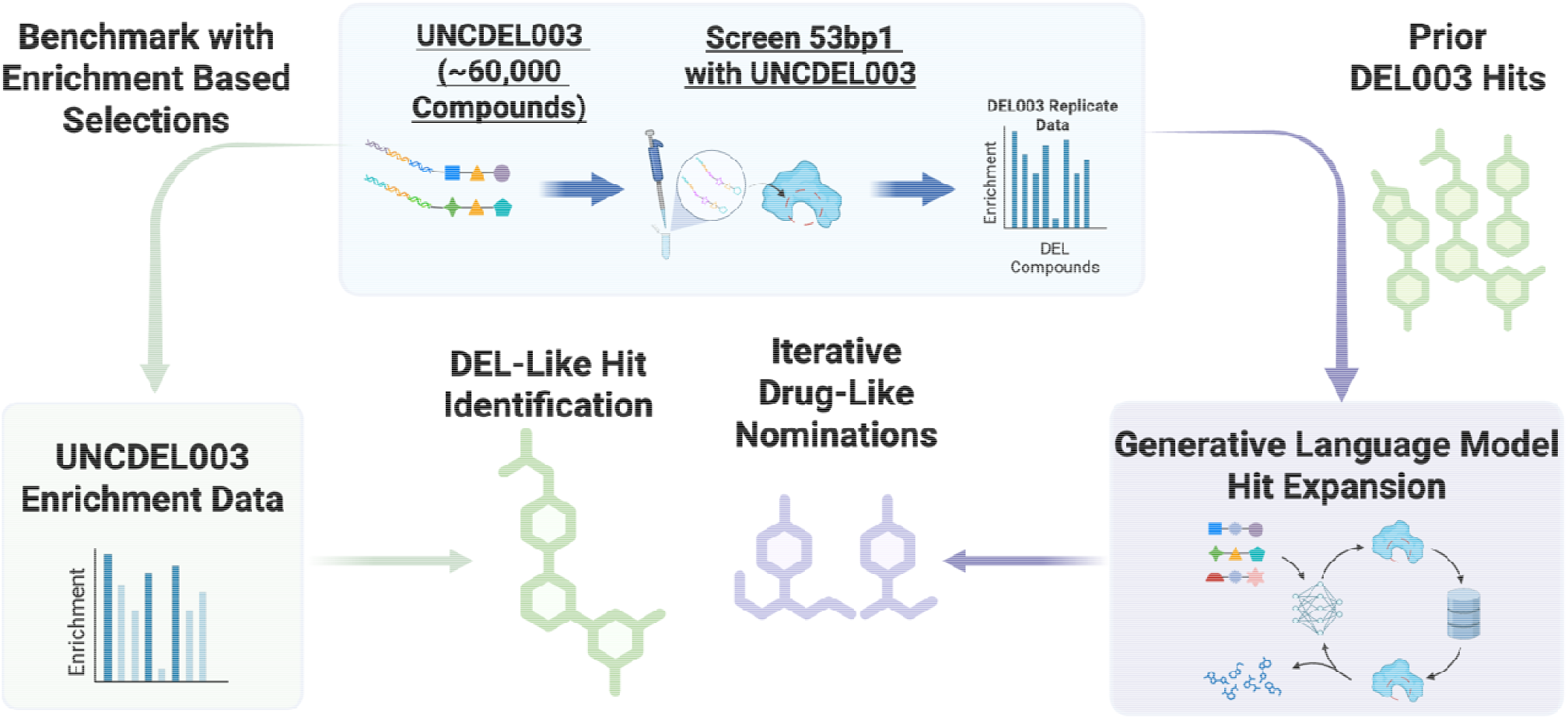
Workflow for the 53BP1 hit-finding campaign. The schematic illustrates a dual-pathway approach comparing traditional enrichment-based selection methods against a generative model pipeline. Prior hits derived from screening the UNCDEL003 library against 53BP1 are used to seed a generative language model (HIDDEN GEM). This model undergoes iterative cycles of hit expansion to produce novel, drug-like nominations from the purchasable Enamine REAL Space. Concurrently, standard selections utilizing UNCDEL003 replicate data identify conventional DEL-like hits.

## RESULTS AND DISCUSSION

### Targeted Screening of 53BP1-TTD Using UNCDEL003 and Enamine Library

The p53 binding protein 1 (53BP1) plays a crucial role in maintaining genomic stability by facilitating DNA damage repair. It promotes non-homologous end joining (NHEJ), thereby limiting the efficiency of homology-directed repair (HDR) in genome editing. Potent small molecule inhibitors of 53BP1 could serve as universal tools to enhance CRISPR-Cas9 gene editing. The protein features a Triple Tudor domain (TTD), which enables recognition of methyl-lysine marks on histones. We utilized the previously reported focused DEL, UNCDEL003, specifically designed for screening Kme reader domains, which has been consistently employed in chemical probe discovery and optimization for these targets.^8,16,25–32^ As we have had prior success in finding hit compounds for 53BP1 from DEL screening,^16,25^ we aimed to expand our hit discovery efforts by leveraging DEL selections to bias generative models. This combination of experimental and virtual screening enables us to cover a larger chemical space and identify novel chemotypes that may not be represented in the physical screening library. To obtain selection data, UNCDEL003 (58,080 compounds) was screened against the 53BP1-TTD as previously reported.^16,25^ To increase confidence in our selection results for training data, we incorporated replicate DEL selections. This approach has been shown to reduce the Poisson-distributed noise inherent to DEL screening, leading to more reliable data and optimized screening conditions.^33,34^

### Hit Identification by Virtual Screening Enamine REAL Space Using a DEL-Initialized HIDDEN GEM Workflow

Once DEL screenings and data processing were completed, we compared two distinct hit identification strategies in a dual pathway approach (**Figure 1**). For the first strategy, we synthesized previously unexplored top compounds based on aggregated enrichment from two in-house UNCDEL003 screenings of 53BP1 (**Table S1**).^25^ We used this as a benchmark for hit discovery via standard DEL analysis methods. For the second strategy, we deployed our HIDDEN GEM workflow^23^. To maintain diversity among purchasable candidates, we first performed a similarity search with our previously discovered UNCDEL003 compound, UNC8531, to the entire Enamine REAL Space.^25^ We then initialized the HIDDEN GEM workflow by using the top 1 million most similar molecules to finetune the generative model. We utilized the structure of the previously reported UNC8531 DEL hit compound co-crystallized with 53BP1 tandem Tudor domains (PDB: 8SWJ) to initiate the structure-based scoring portion of the HIDDEN GEM workflow, following the standard protocol described in the original paper.^25^ This approach integrates a series of iterative molecular dockings, biased generative chemistry, and accelerated similarity searches to identify high-scoring hits from an ultra-large chemical library (see Methods). Starting with 1 million Enamine REAL Space compounds most similar to our DEL hit, two HIDDEN GEM cycles nominated 57 compounds for testing, yielding 14 experimentally validated hit compounds (**Table 1**).

**Table 1.** TR-FRET testing results from the virtual screening of our generative chemistry model revealed novel regions of chemical space within the entire 37 billion compounds of the Enamine REAL “lead-like” library.

| Hidden Gem Cycle 1 |  |  | Hidden Gem Cycle 2 |  |  |
| --- | --- | --- | --- | --- | --- |
| UNC ID | Enamine ID | TR-FRET IC<sub>50</sub> (μM) | UNC ID | Enamine ID | TR-FRET IC<sub>50</sub> (μM) |
| UNC10413788A | Z52064170 | 22 ± 3.6 | UNC10414645A | Z951545060 | 34 ± 3 |
| UNC10413807A | Z31148358 | 43 ± 0.44 | UNC10414641A | Z1941284819 | 58 ± 10 |
| UNC10413800A | Z1033189090 | 54 ± 8.5 | UNC10414653A | Z8721917624 | 61 ± 3 |
| UNC10413811A | Z8185367274 | 58 ± 5.3 | UNC10414658A | Z8721917616 | 68 ± 13 |

\*IC<sub>50</sub> values are the mean ± SD of at least three replicates determined by TR-FRET screening.

### Differences Observed Across Iterative Generations of HIDDEN GEM

To investigate the effects of iterative generation, we selected two sets of confirmed-active generated chemotypes that were conserved across two HIDDEN GEM optimization cycles. The preservation of the same chemotype across successive cycles provided a more interpretable framework for evaluating the medicinal chemistry changes introduced during iterative optimization, allowing us to visualize the structural evolution and property optimization of the four hit compounds relative to the baseline DEL hits (**Figure 2A, Figure 2B**). Comparative analysis of the physicochemical property distributions of HG1 and HG2 nominations revealed statistically significant shifts between cycles in molecular weight (MW; p = 0.0176), heteroatom count (p = 0.0249), and rotatable bonds (p = 0.0304; two-tailed t-test) Furthermore, validated, purchasable hits from both generative cycles exhibited favorable drug-likeness and synthetic accessibility (SA) profiles, demonstrating that iterative generation could diversify chemical space while maintaining desirable molecular properties (**Figure 2B, Figure 2C**).

**Figure 2.**
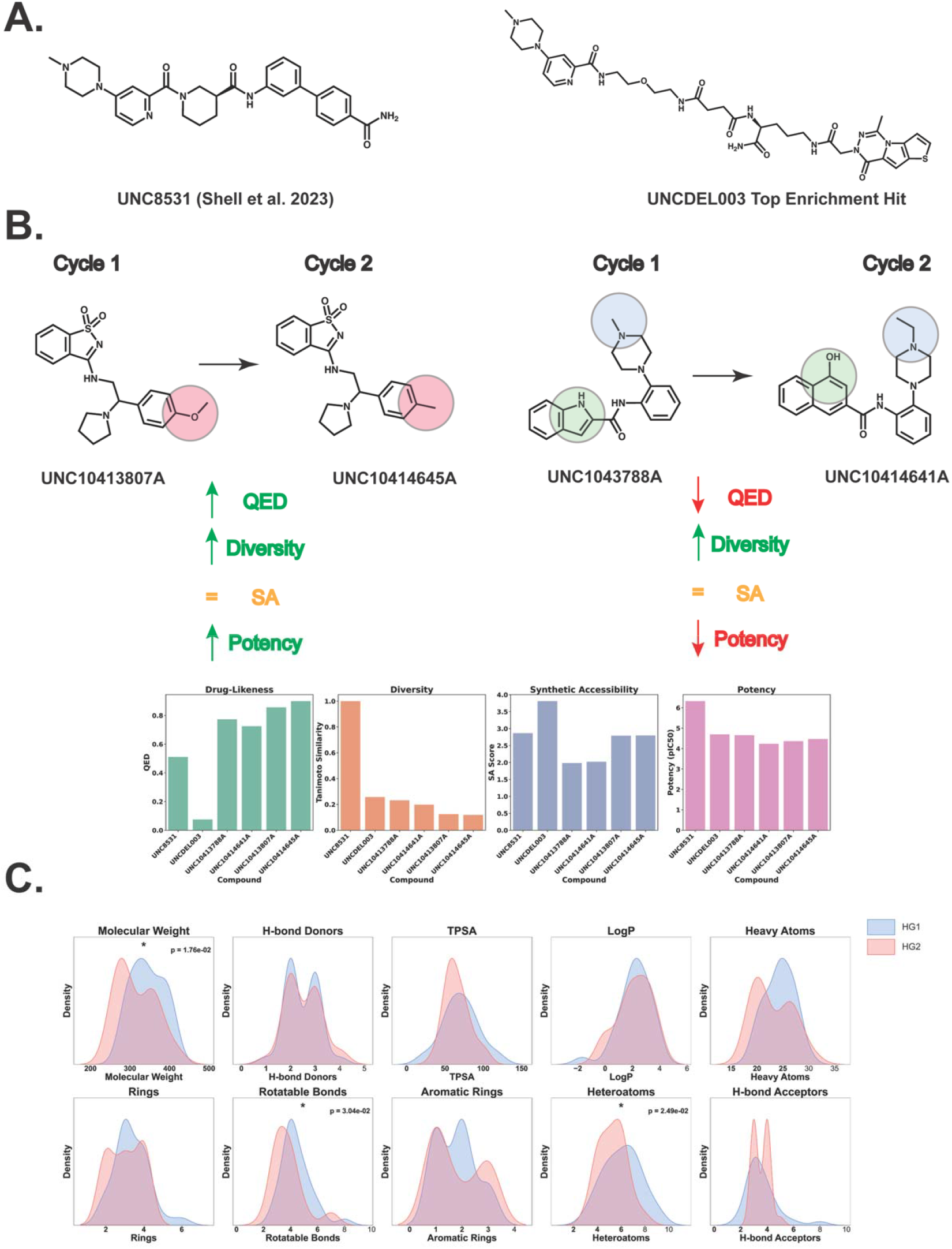
Generative nominations exhibit comparable potency, but superior drug-like properties compared to DEL-based baseline compounds. A) Chemical structures are shown for the previously reported UNC8531 identified through mono and disynthon aggregation, and the top active compound from enrichment-based nominations in the UNCDEL003 screen against 53BP1-TTD. B) Structural evolution and property optimization are detailed for four Enamine REAL hit compounds generated during Cycle 1 and Cycle 2 of HIDDEN GEM, alongside bar charts quantifying drug-likeness (QED), structural diversity (Tanimoto similarity using Morgan fingerprints, 2048 bits, radius 32), synthetic accessibility (SA) scores, and log potency relative to the DEL hits. C) Density distributions of physicochemical properties for all nominations across both generative cycles highlight statistically significant shifts (two-tailed t-test) between cycles in molecular weight, rotatable bonds, and heteroatom count.

While our study experimentally validates the integration of our prior platform encompassing a chemical language model with docking-based scoring, both components are modular. The language model and scoring function could readily be substituted with emerging approaches (e.g., ChemBERTa^35^, Chemformer^36^, Boltz^37^), underscoring the adaptability of HIDDEN GEM to advances in generative chemistry and virtual screening.

### Experimental Validation and Chemical Space Analysis of 53BP1 Hits

To further investigate the distribution shift in chemical space of our HIDDEN GEM hits, we applied non-linear dimensionality reduction using t-distributed stochastic neighbor embedding (t-SNE). In our reduced dimensional space, we can see localization of HIDDEN GEM compounds with unique chemical moieties (**Figure 3A**), such as benzoisothiazole (UNC10413807A), naphthalene (UNC10413811 and UNC10414641), and indole (UNC10414653A) substituents. By combining receptor structural information with generative chemistry^23^, this workflow enabled exploration of distinct regions of chemical space and facilitated the discovery of novel scaffolds not captured by DEL, without the need for extensive medicinal chemistry follow-up or synthon aggregation, as the nominated compounds are readily purchasable and drug-like. The resulting nominations were structurally distinct from the reference compound, with a maximum Tanimoto similarity of 0.23 (cycle 1) and 0.19 (cycle 2), underscoring their novelty (**Figure 3B)**.

**Figure 3.**
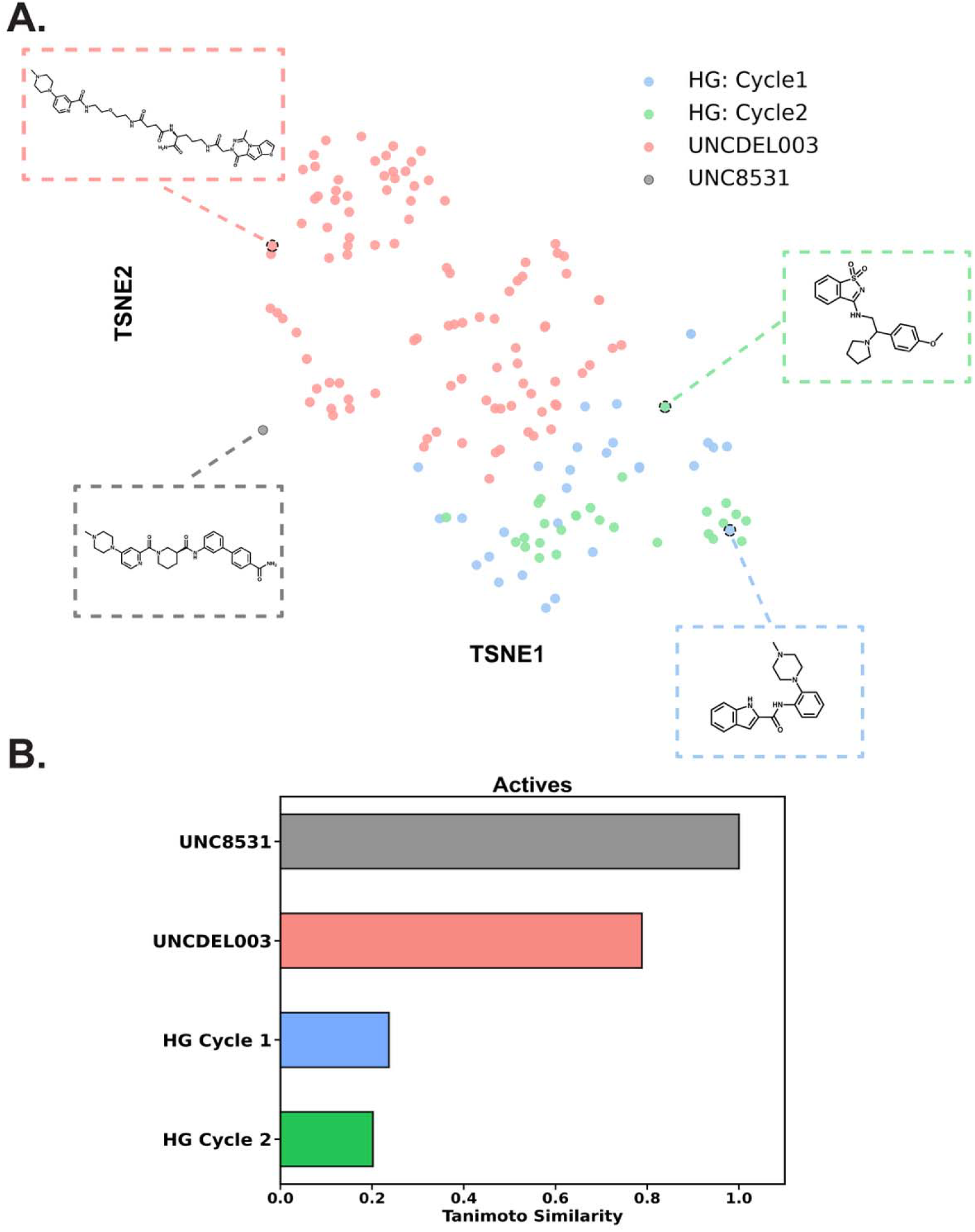
Chemical space mapping and similarity calculations reveal that HIDDEN GEM explores novel regions of chemical space. **A)** t-SNE visualization mapping the chemical space of HIDDEN GEM (HG) Cycle 1 (blue), HG Cycle 2 (green), DEL3 (salmon), and UNC8531 (grey). Representative 2D structures are highlighted for each group. t-SNE analysis was performed using Morgan fingerprints (radius=3, 2048 bits). **B)** Bar chart quantifies the max Tanimoto similarity scores of DEL3, HG Cycle 1, and HG Cycle 2 to the reference compound UNC8531.

### Deep Learning-Based Prediction of Compound Solubility for Assessing Generative Expansion of DEL-Based Hits

We began by experimentally validating the top compound from the DEL screen, UNC10413788A, which showed a TR-FRET IC_50_ of 22 ± 3 μM. Among the most potent compounds, UNC10413788A and two generative chemistry hits exhibited comparable activity against the 53BP1 TTD. Notably, the generative hits extended beyond the chemical space represented by the DEL-derived hit, yielding compounds with greater structural diversity and more drug-like properties (**Figure 4A, Figure 4B**). To further assess their medicinal chemistry profiles, we employed the recently developed FastSolv^38^ deep learning model to predict aqueous solubility. The temperature-dependent solubility profiles predicted by FastSolv indicated that the generative chemistry hits exhibited higher predicted solubility (logS) across the evaluated temperature range (250–350 K) relative to both the baseline control and the traditional DEL-derived hit (**Figure 4C**). These results provide additional evidence that generative expansion can improve drug-like properties beyond conventional computational benchmarks, including synthetic accessibility (SA) and quantitative estimate of drug-likeness (QED) scores.

**Figure 4.**
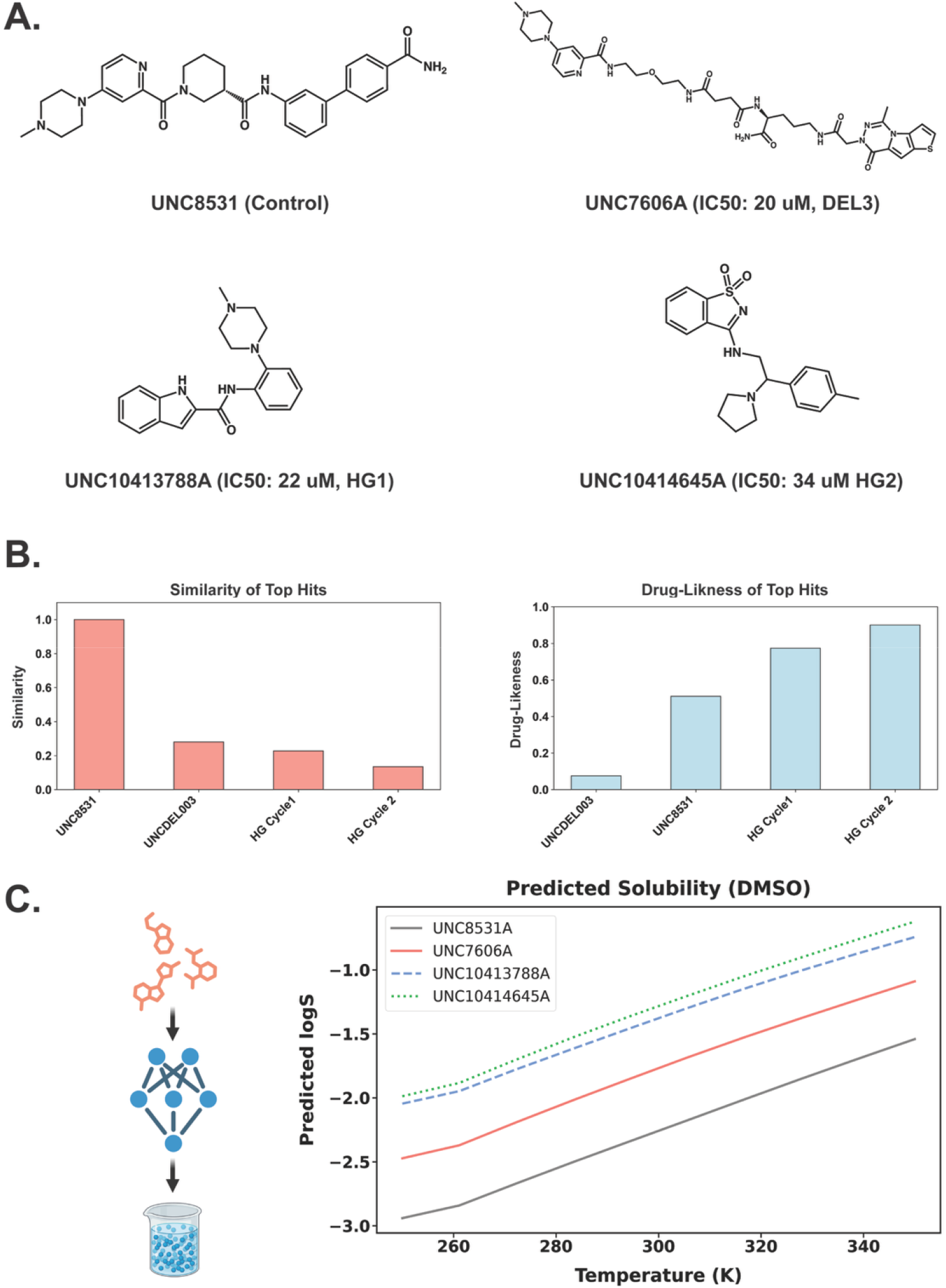
Structural and physicochemical comparison of the most potent hits across discovery methods. **A)** Chemical structures are shown for the original UNCDEL003 tool reference compound, UNC8531 and most potent hits derived from the DEL3 screen (UNC7606A) and generative cycles 1 (UNC10413788A) and 2 (UNC10414645A). **B)** Bar charts quantify the Tanimoto similarity (Morgan fingerprints, radius=3, 2048 bits) of each compound relative to UNC8531 alongside their computed drug-likeness, demonstrating that the generative models (HG Cycle 1 and 2) produce the most chemically distinct active candidates while maintaining superior drug-like properties. **C)** Temperature-dependent solubility profiles in DMSO (logS) were computed for each candidate utilizing FastSolv.^38^ The resulting thermodynamic solubility curves indicate that the generative model nominations exhibit better predicted solubility across a standard temperature gradient (250 K to 350 K) compared to both the baseline control and the traditional DEL hit.

### Structural Modeling of Top Generative Compounds in Complex with 53BP1 TTD

Confirming the binding and interaction mode of this initial hit was critical, as it validated our computational docking model. We stress the importance of validating a docking model, since the HIDDEN GEM workflow relies on its scoring function to nominate all subsequent compounds. With a validated model, HIDDEN GEM successfully identified compounds in novel chemical space. Docking studies confirmed that the top hit compounds retained key molecular interactions similar to the reference compound, including a conserved methyl-piperazine group participating in π-cation interactions (**Figure 5, Figure S1**).

**Figure 5.**
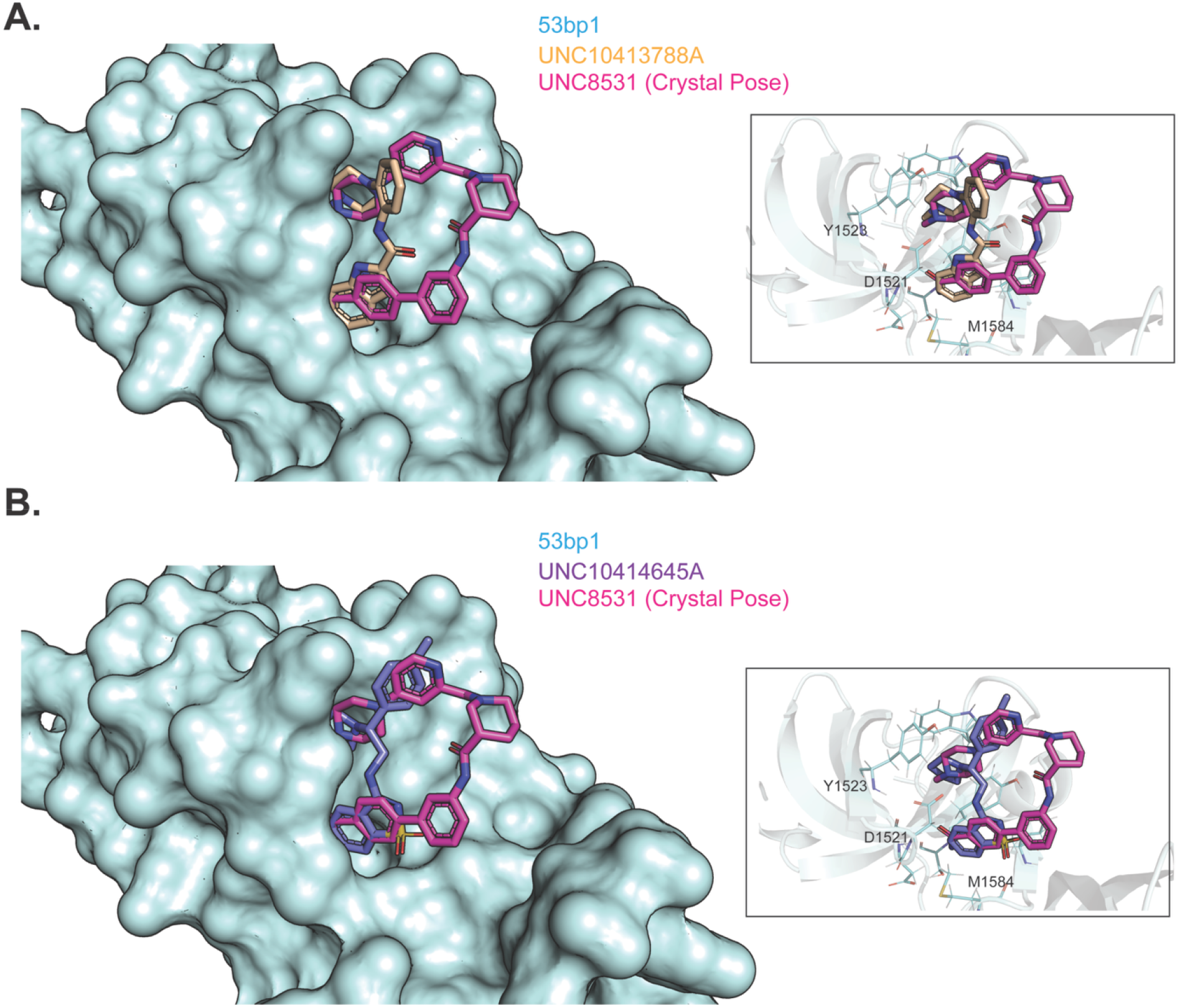
Predicted binding poses and key interactions of top generative chemistry hits UNC10413788A and UNC10414645A. **A)** The top predicted docking pose of UNC10413788A (tan) and **B)** the top docking pose of UNC10414645A (purple) are shown docked within the 53BP1 structure (light blue surface; PDB: 8SWJ). In both panels, the compounds are overlaid with the crystal conformation of the reference ligand UNC8531 (magenta) to illustrate spatial alignment. Detailed inset views for both structures highlight specific binding pocket interactions between the docked models and key residues Y1523, D1521, and M1584.

### Application of HIDDEN GEM to Diversity-Based DELs

While our work demonstrates the application of HIDDEN GEM to a focused DEL, the same workflow is highly valuable for diversity-based DELs, especially those screened through commercial services. Ultra-large, diversity-based DELs are often proprietary platforms managed by Contract Research Organizations (CROs), and a common challenge is that clients typically receive only the structures of a handful of top-binding compounds. This limited data disclosure severely restricts the use of machine learning to explore the chemical space around the initial hits. Our approach directly addresses this bottleneck. By using the few disclosed binders as a starting point, HIDDEN GEM can navigate vast, commercially available chemical libraries to identify novel, purchasable molecules. This strategy effectively expands the search for new scaffolds far beyond the initial proprietary hits, providing a powerful way to maximize the value of any DEL campaign, regardless of data accessibility.

## CONCLUSION

This study demonstrates that an AI-driven, structure-based virtual screening workflow, when combined with limited chemical matter from a focused DNA-encoded library (DEL), can successfully identify novel, potent, and drug-like binders for the 53BP1 tandem Tudor domain (TTD). This integrated strategy expanded the search for hits from a ∼58K-member focused library into the 37-billion-compound Enamine REAL Space (at the time of screening). Our comparative analysis shows that this approach yielded hits with activities comparable to those top enriched members from the original DEL screen, while significantly improving chemical novelty and shifting the distribution toward more favorable drug-like properties.

The resulting nominated compounds from HIDDEN GEM exhibited activities in the low-to mid-micromolar range, greater chemical diversity, better predicted solubility, and superior drug-like properties compared to the initial DEL-derived hits. Their commercial availability further streamlines the hit expansion process. This approach is particularly valuable for smaller or focused DELs, where supervised ML methods can struggle with out-of-domain predictions. Ultimately, the flexible workflow presented here serves as a powerful blueprint for leveraging focused DEL screening data to discover diverse, tractable hits from vast commercial libraries. This synergy between DEL data and generative AI not only represents a significant advance for DEL data analysis but also highlights the value of incorporating such datasets into future open machine learning initiatives to fuel the next generation of drug discovery models.

## Materials and Methods

Reagents, protein expression/purification, DEL production and screening, and TR-FRET screening protocols were previously reported.^16,25^

### DEL Data Preparation for HIDDEN GEM and Experimental Validation

Using two replicates of UNCDEL003 selections against 53BP1, the aggregated enrichment values were first obtained using our open-source DEL informatics software, DELi.^39^ The DELi pipeline processed sequence alignments, applied read-count normalization across the replicates, and computed the final enrichment metrics to account for Poisson-distributed noise. A previously reported hit from UNCDEL003 screening, UNC8531, was selected and used for HIDDEN GEM initialization. Further details on the use of HIDDEN GEM can be found in our previous publication.^23^

### Cheminformatics, Proprty Calculation, and t-SNE Analysis

All compound SMILES string parsing, 2D structure generation, and physicochemical property calculations were executed using RDKit (Python). Chemical similarity was quantified via Tanimoto similarity utilizing Morgan fingerprints (radius=3, 2048 bits). RDKit was also utilized to calculate molecular weight (MW), number of rotatable bonds, heteroatom counts, Quantitative Estimate of Drug-likeness (QED), and Synthetic Accessibility (SA) scores. Non-linear dimensionality reduction of the chemical space was performed using t-distributed stochastic neighbor embedding (t-SNE) implemented in scikit-learn, utilizing the computed Morgan fingerprints with a perplexity of 30, a learning rate of 200, and 1000 iterations.

### FastSolv Predictive Solubility Modeling

Temperature-dependent thermodynamic solubility (logS) profiles were predicted using FastSolv,^38^ a deep learning-based property prediction model. Computations were accelerated using CUDA parallel computing infrastructure. Solubility predictions were executed for the selected candidates across a simulated temperature gradient spanning from 250 K to 350 K in DMSO to assess structural stability and formulation viability.

### HIDDEN GEM Virtual Screening and Nomination

The HIDDEN GEM workflow was executed following the standard protocol described in the original publication. Briefly, the virtual screening process begins with an initialization step, in which a relatively small compound library is docked into 53BP1 using Glide (see Glide methods below). In this study, we used a set of 1 million Enamine REAL Space compounds most similar to UNC8531, rather than the diverse initialization set described in the original HIDDEN GEM paper. The top 1K compounds, ranked by Glide docking scores, were then used to bias a ChEMBL-pretrained generative model released with the HIDDEN GEM publication^23^, which generated 100K de novo small molecules predicted to have favorable binding scores against 53BP1. This new set was subsequently redocked, and the top 1K scoring molecules were used as queries for a large-scale chemical similarity search. This search identified up to 10K commercially available compounds from Enamine REAL Space that were highly similar to the generated queries. These candidate molecules were then docked and scored, and the top 1K were selected for visual inspection and final compound selection for purchase. Additionally, these top candidates were used to further bias the generative model in a second cycle of the HIDDEN GEM workflow.

As demonstrated in the initial publication and in the current work, this iterative approach that combines initial docking, generative chemistry, similarity-based expansion, and final docking, enabled HIDDEN GEM to efficiently converge on chemical subspace diverse from the initial DEL hit.

### Glide Docking and Receptor Preparation

Docking experiments were conducted using the UNC8531-bound form of the 53BP1 tandem Tudor domain (PDB ID: 8SWJ). Prior to docking, the structure of the protein was prepared using Schrödinger Protein preparation tools with default settings: adding and optimizing hydrogen atoms, setting residue ionization and tautomer states with PROPKA at pH 7, removing water molecules more than 3 Åaway from the HET atoms, and performing restrained structure minimization with the OPLS4 force field to converge the heavy atoms to an RMSD of 0.3 Å.^40^ This offers minor structure optimization to ensure optimal performance of subsequent docking runs. Following protein preparation, the Glide receptor grid was generated, with the grid box centered on UNC8531 ligand and the inner and outer box sizes were set to 10 Å and 20 Å, respectively. The structural model was validated by assessing the docked pose for UNC8531, which was found to have <1.0 Å RMSD from the initial crystal ligand pose. The structures of all ligands were prepared using the LigPrep tool to sample up to 32 conformation states during the docking protocol.^41,42^ Glide Standard Precision (SP) docking was then conducted for the series of ligands prepared in the previous step. Ligands were ranked according to their Glide SP docking scores, using the best-scoring pose for each compound as its representative.^43^

### General Method for Compound Synthesis

Selected hit molecule syntheses were obtained by solid-phase peptide synthesis techniques on Fmoc-Rink Amide MBHA resin. Compound syntheses were performed in a polypropylene syringe fitted with a porous polyethylene frit. Solvents and soluble reagents were removed by applying pressure on the syringe piston. NMP can be substituted for DMF in this protocol. Fmoc-Rink Amide MBHA (0.01 mmol) resin was soaked in 1 mL of DCM for 30 min on a tabletop shaker. After removing the solvent, resin was soaked in NMP solvent (1 mL) and allowed to shake for 15 min. Solvent was drained, and the syringe was filled with a 25% piperidine (or 2.5% pyrrolidine and DBU) solution in NMP (1 mL) and allowed to shake for 10 min. The deprotection solution was drained, and the resin was washed thrice with NMP solvent (3 x 2 mL). Then, the pre-activated Fmoc-amino acid (Fmoc-AA) in NMP was loaded into the syringe and shaken for 1 hour. [Pre-activation of Fmoc-AA: To the Fmoc-AA (0.03 mmol) dissolved in NMP (1 mL), DIPEA (0.05 mmol) was added and followed by HATU coupling reagent (0.03 mmol) in NMP (1 mL) and allowed to sit for 10 min.] Washings between deprotection, coupling, and subsequent deprotection steps were carried out with NMP or DMF (3 × 1 min) using 2 mL of solvent per 0.2 mmol of resin for each wash. After completion of the compound sequence, the resin was washed with NMP (2 x 3 mL) followed by DCM (2 x 3 mL). Then, the syringe was filled with a cleavage cocktail (1 mL) and shaken for 1 hour. [Cleavage cocktail preparation: TFA:H_2_O:TIPS-95:2.5:2.5 respectively.] Then, the TFA solution was collected in a glass vial and evaporated. The crude compound was purified using an Agilent Technologies 1260 Infinity preparative-HPLC [Solvent system: water:acetonitrile (90:10); Flow rate: 40 mL/min; UV detector wavelength: 254 nm] to obtain a pure final product. Final compounds were verified by LCMS prior to testing.

Analytical LCMS data for compounds were acquired on an Agilent 6110 Series system with UV detector set to 220 nm, 254 nm, and 280 nm. Samples were injected onto an Agilent ZORBAX Eclipse Plus 4.6 X 50 mm, 1.8 µm, C18 column at 25°C. Mobile phases A (H_2_O + 0.1% acetic acid) and B (ACN + 1% water and 0.1% acetic acid) were used in a linear gradient from 10% to 100% B in 5 min, followed by a flush at 100% B for another 2 min with a flow rate of 1.0 mL/min. Mass spectra (MS) data were acquired in positive ion mode using an Agilent 6110 single quadrupole mass spectrometer with an electrospray ionization (ESI) source.

### Chemistry

The general chemistry scheme for all synthesized compounds is provided in the Supporting Information. All compounds are >95% pure by HPLC analysis.

## ASSOCIATED CONTENT

### Supporting Information

**Figure S1**: Chemical Similarity Evaluation from Each Method; **Figure S2**: TR-FRET Data from HG Hits; **Figure S3**: UNCDEL003 Compound Synthesis and Reagents; **Table S1**: Top 10 DEL Compounds Based on Aggregated Compound Frequency

## AUTHOR INFORMATION

### Notes

The authors declare no competing financial interest.

## ACKNOWLEDGMENT

The authors thank Ivanna Zhilinskaya for reviewing the experimental data and Jacqueline L. Norris-Drouin for protein production (as referenced in previous work). The authors gratefully acknowledge support from the NIH Biophysics Training Grant (T32GM148376-01A1). We also thank Dr. Caroline Daigle, Dr. Stephen Frye, and Dr. Lindsey James for helpful discussions and suggestions during this research.

## ABBREVIATIONS

53BP1: p53 binding protein 1
AA: amino acid
ACN: acetonitrile
AI: artificial intelligence
DBU: 1,8-diazabicyclo[5.4.0]undec-7-ene
DCM: dichloromethane
DEL: DNA-encoded library
DIPEA: N,N-diisopropylethylamine
ESI: electrospray ionization
FCFP: functional class fingerprints
HATU: hexafluorophosphate azabenzotriazole tetramethyl uronium
HG: HIDDEN GEM
HPLC: high-performance liquid chromatography
HTS: high throughput screening
Kme: methyl-lysine
MBHA: 4-methylbenzhydrylamine
ML: machine learning
MS: mass spectra
NGS: next generation sequencing
NMP: N-methylpyrrolidone
TFA: trifluoroacetic acid
TIPS: triisopropyl silane
TR-FRET: time-resolved fluorescence resonance energy transfer
TTD: tandem Tudor domain.

## DATA AND SOFTWARE AVAILABILITY

The authors present the Compound IDs, SMILES, and associated assay activity of all compounds tested in the manuscript within the Supplemental Materials. In addition, all compound SMILES and aggregated enrichment are available within the Supplemental Materials.

---

**Figure S1.**
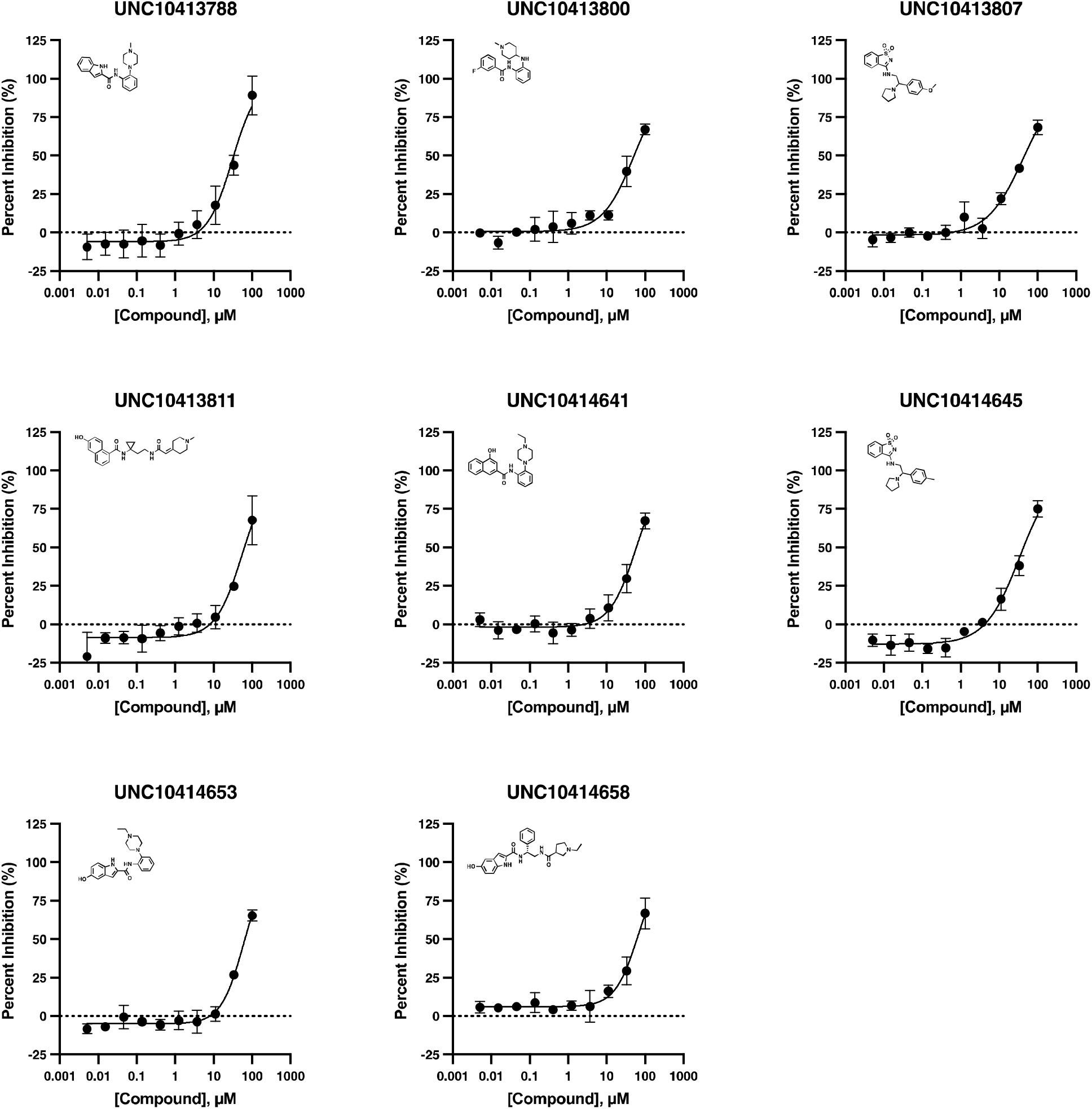
53BP1 TR-FRET displacement assay dose-response curves for compounds identified from HIDDEN GEM. Each data point represents the mean of three replicates (n=3), with error bars indicating standard deviation. Dose-response curves were fitted using a four-parameter Hill equation.

**Figure S2.**
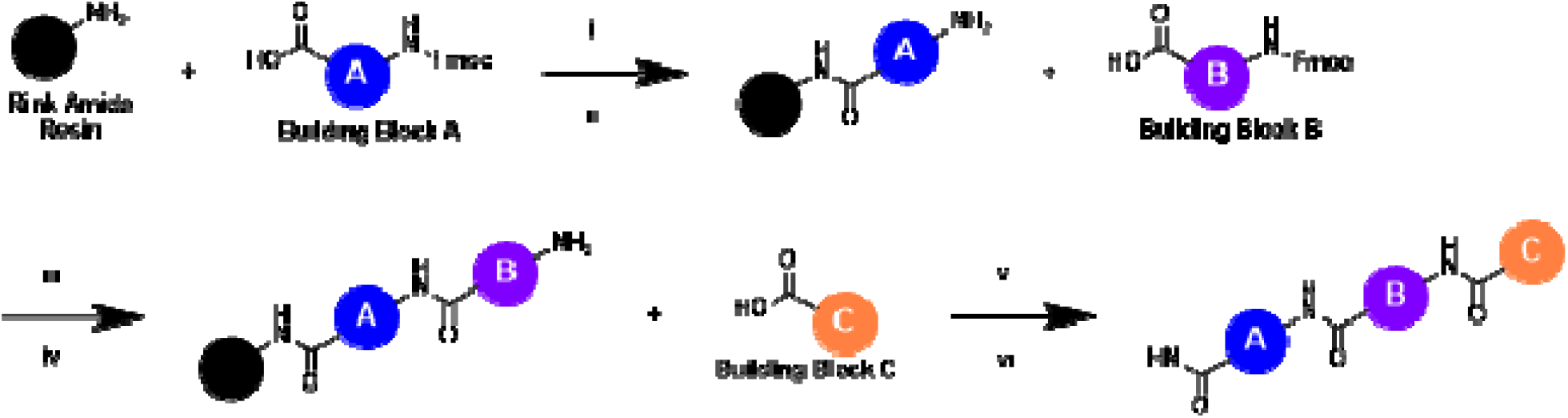
UNCDEL003 Compound Synthesis. **Reagents and conditions**: i) Solid phase synthesis on Rink Amide resin, 3’-((((9H-fluoren-9-yl)methoxy)carbonyl)amino)-[1,1’-biphenyl]-4-carboxylic acid, HATU, DIPEA, NMP or DMF, rt; ii) Fmoc deprotection, 25% piperidine in NMP, rt; iii) Amide coupling with building block B, HATU, DIPEA, NMP or DMF, rt; iv) Fmoc deprotection, 25% piperidine in NMP, rt; v) Amide coupling with building block C, HATU, DIPEA, NMP or DMF, rt; vi) Cleavage from Rink Amide resin, 95% v/v TFA, 2.5% v/v TIPS, and 2.5% v/v H_2_O, rt

**Table S1.** Top 10 DEL Compounds Based on Aggregated Compound Frequency.

| **A Synthon** | **B Synthon** | **C Synthon** | **Compound ID** | **Aggregated**<br>**Compound**<br>**Frequency** |
| --- | --- | --- | --- | --- |
| A12 | B22 | C6 | A12-B22-C6 | 82 |
| A10 | B22 | C6 | A10-B22-C6 | 74 |
| A8 | B22 | C6 | A8-B22-C6 | 66 |
| A30 | B25 | C6 | A30-B25-C6 | 62 |
| A6 | B20 | C6 | A6-B20-C6 | 61 |
| A2 | B20 | C6 | A2-B20-C6 | 60 |
| A7 | B22 | C6 | A7-B22-C6 | 56 |
| A26 | B39 | C6 | A26-B39-C6 | 55 |
| A26 | B28 | C6 | A26-B28-C6 | 54 |
| A14 | B22 | C6 | A14-B22-C6 | 53 |

